# Wnt ligands are not required for planar cell polarity in the *Drosophila* wing or notum

**DOI:** 10.1101/2020.06.05.137182

**Authors:** Ben Ewen-Campen, Typhaine Comyn, Eric Vogt, Norbert Perrimon

**Author notes:** Corresponding Author: Norbert Perrimon, Harvard Medical School, New Research Bldg., Room 336G, Howard Hughes Medical Institute, 77 Ave. Louis Pasteur, Boston, MA 02115.

## Abstract

The *frizzled* (*fz*) and *disheveled* (*dsh*) genes are highly conserved members of the core planar cell polarity (PCP) pathway and of the Wnt signaling pathway. Given these dual functions, a number of studies have examined whether Wnt ligands may provide a global, tissue-scale orientation cue for PCP establishment during development, and these studies have reached differing conclusions. In this study, we re-examine this issue in the *Drosophila melanogaster* wing and notum using split-Gal4 co-expression analysis, systematic pairwise and triple somatic CRISPR-based knock-outs and double RNAi experiments. Pairwise loss-of-function experiments targeting *wg* together with other Wnt genes does not produce PCP defects, neither via somatic CRISPR nor RNAi. In addition, somatic knock-out of *evi* (aka *wntless*), which is required for the secretion of all Wnt ligands expressed in these tissues, did not produce detectable PCP phenotypes. Altogether, we were unable to find support for the hypothesis that Wnt ligands contribute to PCP signaling in the *Drosophila* wing or notum.

## Introduction

Planar cell polarity (PCP) refers to the coherent orientation of cells in a sheet, and is driven by a conserved molecular pathway that is required for the proper development and function of many animal tissues (Adler, 2012; Butler and Wallingford, 2017; Goodrich and Strutt, 2011; Yang and Mlodzik, 2015a). Decades of functional analyses in the epithelia of *Drosophila* and other model organisms have led to the identification of six highly conserved “core” pathway members, whose asymmetric localization and function within cells drives downstream manifestations of PCP, both autonomously and non-autonomously. In cells of the developing fruit fly wing, the transmembrane protein Frizzled (Fz) and the cytosolic proteins Disheveled (Dsh) and Diego (Dgo) accumulate at the distal cortex. At the proximal cortex, an asymmetric signaling complex forms that includes the transmembrane protein Strabismus aka Van Gogh (Stbm/Vang) and the cytosolic protein Prickle (pk). The sixth core pathway member, Flamingo aka Starry night (Fmi/Stan) accumulates both proximally and distally (Adler, 2012; Butler and Wallingford, 2017; Goodrich and Strutt, 2011; Yang and Mlodzik, 2015). Loss-of-function of any core pathway members leads to PCP defects, which ultimately manifest in mispolarization of the actin-rich wing hairs (Gubb and García-Bellido, 1982; Wong and Adler, 1993). In addition to the core pathway, a second PCP pathway that includes Fat, Daschous, and Four-jointed has been identified in *Drosophila* (Adler, 2012; Butler and Wallingford, 2017; Goodrich and Strutt, 2011; Yang and Mlodzik, 2015a).

At relatively small spatial scales, feedback interactions amongst core PCP pathway components provide a working molecular model for PCP within cells and their neighbors. However, there remains active debate regarding the molecular mechanisms that allow core pathway members to correctly orient with regard to “global” patterning at the level of whole tissues. Researchers have long hypothesized there may be a secreted molecule (referred to as “Factor X” by (Lawrence et al., 2002)) that coordinates tissue-scale morphogenesis to the PCP pathway. Among the possible candidates for “Factor X”, Wnt ligands have long been considered “obvious candidates for such factors” (Goodrich and Strutt, 2011) because both Fz and Dsh are highly conserved members of this signaling pathway: Fz and its paralog Fz2 function as co-receptors for Wnt ligands (Bhanot et al., 1999; Chen and Struhl, 1999; Müller et al., 1999), and Dsh is also a highly conserved component of the Wnt ligand-receptor signalosome (Perrimon and Mahowald, 1987; Sharma et al., 2018). However, a number of loss-of-function studies of Wnt ligands have failed to find any PCP phenotypes (Chen et al., 2008; Lawrence et al., 2002). Specifically, (Lawrence et al., 2002) examined clones in the adult abdominal epithelium homozygous for a deletion covering four of the seven *Drosophila* Wnt ligands (*wg, wnt4, wnt6*, and *wnt10*), and found no PCP defects within or nearby such clones. (Chen et al., 2008) generated quintuple clones in the wing lacking *wg, wnt2, wnt4, wnt6*, and *wnt10*, as well as clones lacking *porcupine*, a gene necessary for the correct lipid modification of Wnt ligands, and found no evidence of PCP defects in either case. In addition, homozygous mutants for *evi* (aka *wntless*), which are defective in Wnt secretion, display wildtype PCP (Bartscherer et al., 2006).

Because Wg is required in early larval stages to specify the presumptive wing (Ng et al., 1996), the above studies relied on the analysis of *wg* mutant clones in an otherwise wildtype background, and thus could not conclusively rule out a role for long-range Wnt signaling from surrounding tissues. In contrast, gain-of-function studies have demonstrated that over-expression of either Wg or Wnt4 (but not other Wnt ligands) in ectopic regions of the developing wing can re-orient the polarity of surrounding wing hairs (Lim et al., 2005; Wu et al., 2013), and that this reorientation requires *fz* (Wu et al., 2013). In addition, a study that bypassed the early requirements for *wg* using a temperature sensitive mutation indicated that double mutant flies lacking *wg* and *wnt4* displayed statistically significant PCP phenotypes in the pupal and adult wing (Wu et al., 2013). Furthermore, studies in vertebrates have implicated certain Wnt ligands in PCP signaling (reviewed in Yang and Mlodzik, 2015). Taken together, the evidence on whether Wnts, either individually or in combination, influence PCP signaling remains inconclusive and actively debated.

In this study, we re-examine whether Wnt ligands are required for PCP using systematic double knock-outs or knock-downs via Gal4-UAS driven somatic CRISPR or RNAi, respectively. We independently analyzed the expression of each Wnt using a new collection of knock-in split-Gal4 reporters, and then performed multiplex somatic CRISPR using short guide RNAs (sgRNAs) targeting *wg* in combination with each of the other six Wnt genes, as well as various combinations thereof. In addition, we use somatic CRISPR to target *evi* (*wntless*), a protein required for the secretion of Wnt genes except *wntD* (Herr and Basler, 2012). Lastly, we perform double RNAi experiments against *wg, wnt4*, and *wnt6* in the wing and in the notum. We did not detect PCP phenotypes in any of these knock-out or knock-down conditions, in either wing or notum. Our results suggest that Wnts do not impinge on PCP signaling. We note that our data are consistent with a study recently posted on Biorxiv (Yu et al., 2020), which we discuss below.

## Results

### Split-Gal4 knock-in reporters confirm that multiple Wnt genes are co-expressed in the larval wing disc

To characterize which Wnt genes are expressed in the wing disc during the period where PCP patterning is established in the developing wing, we systematically re-examined the expression and co-expression of the seven *Drosophila* Wnt family genes. In the wing disc, core PCP components progressively acquire asymmetrically localization during pupal stages, and there is also evidence that at least some polarity information exists during L3 larval stages in the form of coherent Fmi/Stan localization that is subsequently remodeled during pupal stages (Classen et al., 2005).

To independently examine which Wnt ligands are expressed during wing development, we used CRISPR-based homology directed repair to knock in split-Gal4 reporter cassettes into an early exon of each Wnt gene (**Figure 1A**) (Gratz et al., 2014; Pfeiffer et al., 2010). These cassettes include an in-frame T2A self-cleaving peptide, followed by either the p65 activation domain or the Gal4 DNA binding domain (DBD), and thus allow for intersectional labeling of cells expressing multiple Wnt genes. Pilot experiments with *wg* and *wnt2* indicated that this approach recapitulates previously described endogenous gene expression of *wg* and *wnt2*, for both p65 and Gal4DBD knock-ins (**Figure S1**), although we note that our *wnt2* knock-ins drove expression in the notum, but not in the wing pouch as has been reported by others (Chen et al., 2008; Kozopas and Nusse, 2002). As predicted, homozygous *wg* knock-ins were lethal, and *wnt2* knock-in homozygotes were male sterile and displayed characteristic wing posture abnormalities (Kozopas and Nusse, 2002; Kozopas et al., 1998). We ultimately created split-Gal4 knock-in reagents for each of the seven Wnt genes.

**Fig 1.**
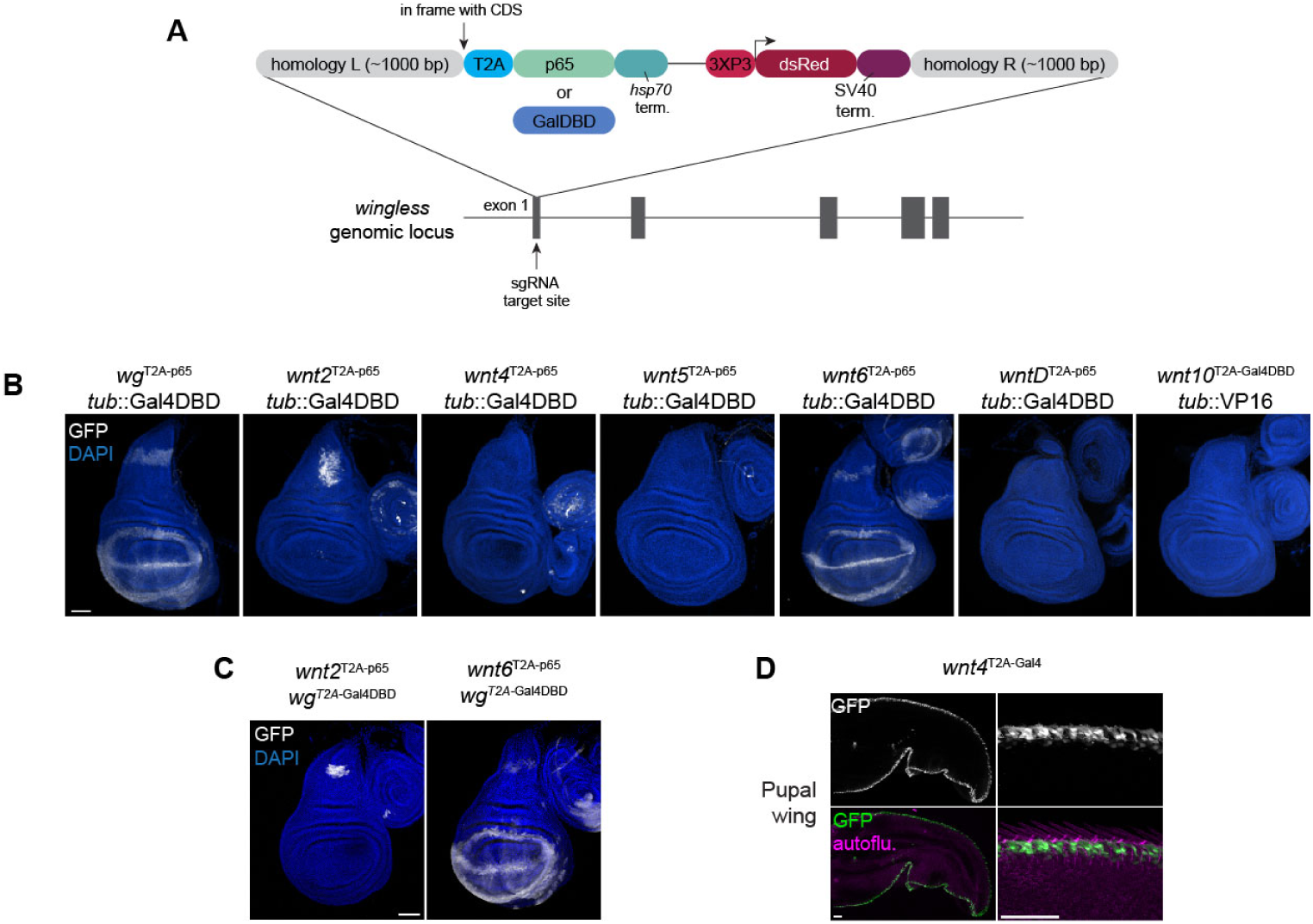
Analysis of Wnt gene expression and co-expression using split-Gal4 reporters. (A) Schematic of knock-in constructs used to generate in-frame split-Gal4 reporters in each of the seven Wnt ligands. (B) Expression pattern of each reporter construct, visualized by crossing to a ubiquitously expressed reciprocal split-Gal4 reporter, visualized with UAS:EGFP or UAS:EYFP (see Methods for details). (C) Intersection of of *wg* expression domain with *wnt2* and *wnt6*. (D) *wnt4* expression in the wing margin of a pupal wing, visualized with an independent Gal4 knock-in driving UAS:GFP. Right-hand images show a magnification of the developing wing margin. Scale bars = 50 µm.

We crossed this collection of split-Gal4 reagents to a ubiquitously-expressed cognate (Gal4DBD or VP16 driven by the *tubulin* promoter) and examined the expression in larval wing discs via a UAS-driven fluorophore (**Figure 1B**). We observed expression of *wg, wnt2*, and *wnt6* in the L3 wing disc, largely consistent with previous reports (Baker, 1988; Janson et al., 2001; Kozopas and Nusse, 2002). Split-Gal4 intersection analysis confirmed that *wg* and *wnt2* are coexpressed in the epithelium of the developing notum, and that *wg* and *wnt6* are extensively co-expressed throughout their entire expression domain at this stage (Janson et al., 2001) (**Figure 1C**). We further characterized these split-Gal4 lines in various adult stem cell populations of the ovary and gut, and in the larval CNS, which we summarize in **Supplementary Note 1** and **Figures S3 and S4**.

Unexpectedly, while we did observe *wnt4* expression in the margin of the pupal wing (**Figure 1D**), we did not detect *wnt4* expression in the dorsoventral (DV) boundary at the L3 larval stage, as has been demonstrated via RNA *in situ* hybridization (Chen et al., 2008; Gieseler et al., 2001; Harris et al., 2016) and a GFP knock-in (Yu et al., 2020). Instead, our *wnt4* reporter was exclusively expressed in a small cluster of cells at the anterior ventral boundary of the wing pouch (**Figure 1B, S2**). Given this discrepancy between our reporter line and previous studies, we examined the expression pattern of an existing knock-in Gal4 reporter from the CRIMIC collection, which contains a “trojan exon” including a splice acceptor and T2A-Gal4, inserted in the first intron of *wnt4* (Lee et al., 2018). The expression of *wnt4*^*CRIMIC*^ was consistent with our split-Gal4 reporter in the L3 wing disc, and largely consistent in the larval CNS (**Figure S2**). As an additional test, we generated a third knock-in reporter (using T2A-Gal4) using an independently-derived donor plasmid (Bosch et al., 2019), inserted into the final exon of *wnt4*. This independent Gal4 insertion (*wnt4*^*T2A-Gal4*^) was consistent with the other two tested in this study, and did not show detectable expression in the DV boundary at the L3 stage (**Figure S2**). To test whether *wnt4* may be expressed in the wing disc during earlier stages, we crossed *wnt4*^CRIMIC^ line to G-TRACE, a lineage-tracing tool that reports both current and past Gal4 expression using two different fluorophores (Evans et al., 2009). This analysis did not reveal any past expression in the wing disc aside from the small cluster of cells on the border of the wing pouch. Importantly, however, we did detect *wnt4* in the wing margin during pupal stages (**Figure 1D**), which has also been described by (Yu et al., 2020), and which is consistent with a possible role in PCP patterning.

### Systematic pairwise knock-outs of Wnt genes using somatic CRISPR

Previous studies in *Drosophila* have demonstrated that ectopic expression of either Wg or Wnt4 can re-orient PCP patterning in surrounding wing hairs (Lim et al., 2005; Wu et al., 2013), and Wu *et al*. (2013) have reported that double mutants for *wg* and *wnt4* display disrupted PCP signaling in the pupal and adult wing. We sought to systematically address the role of Wnt ligands in PCP signaling using *in vivo* somatic CRISPR double- and triple-knockouts. For these experiments, we identified two effective sgRNAs to target each Wnt ligand (see Methods), and created transgenic lines expressing sgRNAs targeting either one or two Wnt genes simultaneously (with either two or four sgRNAs total, respectively) under UAS control. We confirmed that these sgRNAs cleaved their intended target *in vivo* both for single- and double-knockout experiments (**Figure S5**).

To avoid earlier roles for Wg, we used the *nubbin-*Gal4 driver to express both Cas9 and sgRNAs. *nubbin-*Gal4 is expressed throughout the wing pouch beginning in late L2 larval stages (Zirin and Mann, 2007), and thus this approach allowed us to permanently remove target gene function throughout L3, pupal, and adult stages. Control experiments targeting *fz* confirmed that this approach produces strong PCP phenotypes in the adult wing (**Figure S6**), and that *frizzled* is unique among the four Fz-family genes in producing this phenotype (**Figure S6**). Although we did not detect expression of reporters for *wnt5, wntD* or *wnt10* in the wing disc, we included them in our functional studies in order to rule out the possibility that one or more of these ligands is expressed at low levels.

We produced somatic knockouts of *wg* together with each of the other six Wnt genes, and screened for defects in both wing morphology and PCP. As expected, the wing margin was disrupted in all pairwise knock-outs that included *wg* (**Figure 2**). However, in each case, wing hair orientation was indistinguishable from wildtype (see **Figure S6** for examples of PCP phenotypes caused by disrupting *fz*). We next tested pairwise combinations of *wnt4, wnt6*, and *wnt10*, which are part of a genomic cluster on the second chromosome, as well as a triple knockout of *wg, wnt4*, and *wnt6*. We did not observe disrupted PCP in any of these conditions (**Figure 2**). Lastly, we used this approach to knock out *evi* (*wntless*), which is required for the secretion of all Wnt genes except WntD (Bänziger et al., 2006; Bartscherer et al., 2006; Goodman et al., 2006; Herr and Basler, 2012). Knocking out *evi* led to loss of wing margin, essentially phenocopying *wg* loss-of-function, and did not produce PCP defects, consistent with observations from *evi* mutant pharate adults (Bartscherer et al., 2006) (**Figure 2**). Together, these results do not support the hypothesis that Wnt ligands direct PCP orientation in the *Drosophila* wing.

**Fig 2.**
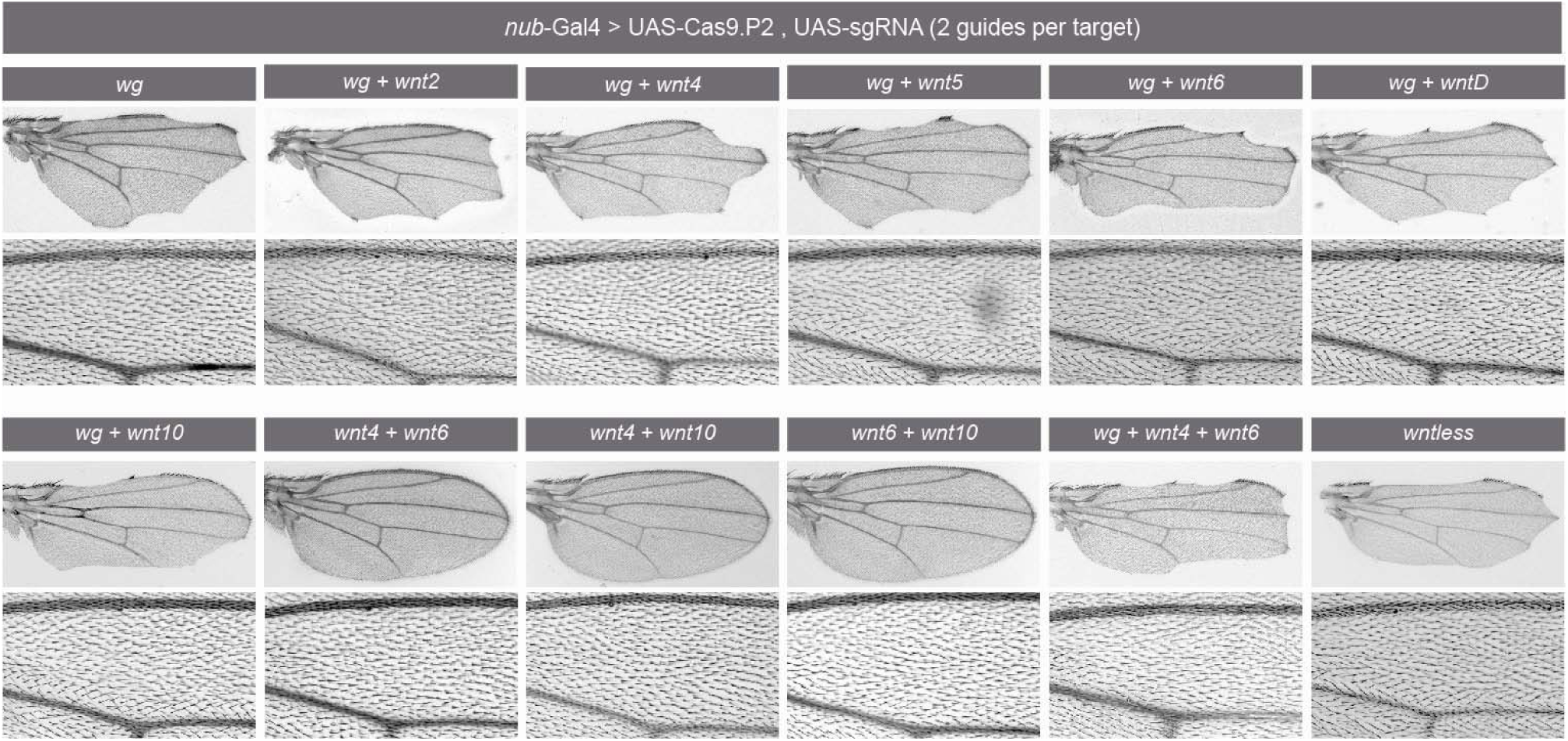
Multiplexed somatic CRISPR knock-out of Wnt genes or *wntless* does not disrupt planar cell polarity. sgRNAs targeting the indicated genes (two guides per target gene) were co-expressed with Cas9.P2 in the developing wing using *nub-Gal4*. Overall wing morphology is shown in the top rows, and magnified views of wing hair orientation in the L3-L4 intervein region are shown in bottom rows. Images show representative images of approximately 20 wings analyzed per genotype.

### Pairwise RNAi against *wg* together with *wnt4* or *wnt6* does not produce PCP phenotypes in the wing or notum

Somatic Gal4-UAS-based CRISPR has been shown to be a powerful and scalable new technique for *in vivo* knockouts (Port and Bullock, 2016; Port et al., 2020). However, one inherent drawback of somatic CRISPR is that it creates mosaic tissues containing both mutant and wildtype cells, and does so variably between individuals. This can be seen, for example, in the variable phenotypes and incomplete loss of margin tissue observed when targeting *wg* (**Figure 2**). In the case of morphogens such as Wnt ligands, it is conceivable that a small number of wildtype cells could still be sufficient to produce an instructive cue for PCP orientation.

To address this issue, we performed double RNAi experiments against *wg, wnt4*, and *wnt6* in the developing wing using *nub-*Gal4, using transgenic dsRNA lines recently shown to produce strong loss-of-function phenotypes, and co-expressed with UAS:*dcr2* to strengthen RNAi-mediated knockdown (Barrio and Milán, 2020). Under these conditions, *wg* RNAi led to both a complete loss of the wing margin and a profound growth defect, which was visually similar to the reported effects of inducing apoptosis in *wg*-expressing cells by over-expressing the pro-apoptotic genes *hid* and *rpr* (Yu et al., 2020). However, even in these severely mispatterned wings, PCP appeared wildtype. Similarly, no PCP defects were observed upon pairwise RNAi against *wg* and *wnt4* or *wg* and *wnt6* (**Figure 3**). Incidentally, our results confirmed recent observations that *wnt4* and *wnt6* have opposing effects on modulating the effects of *wg-*dependent growth (Barrio and Milán, 2020).

**Fig 3.**
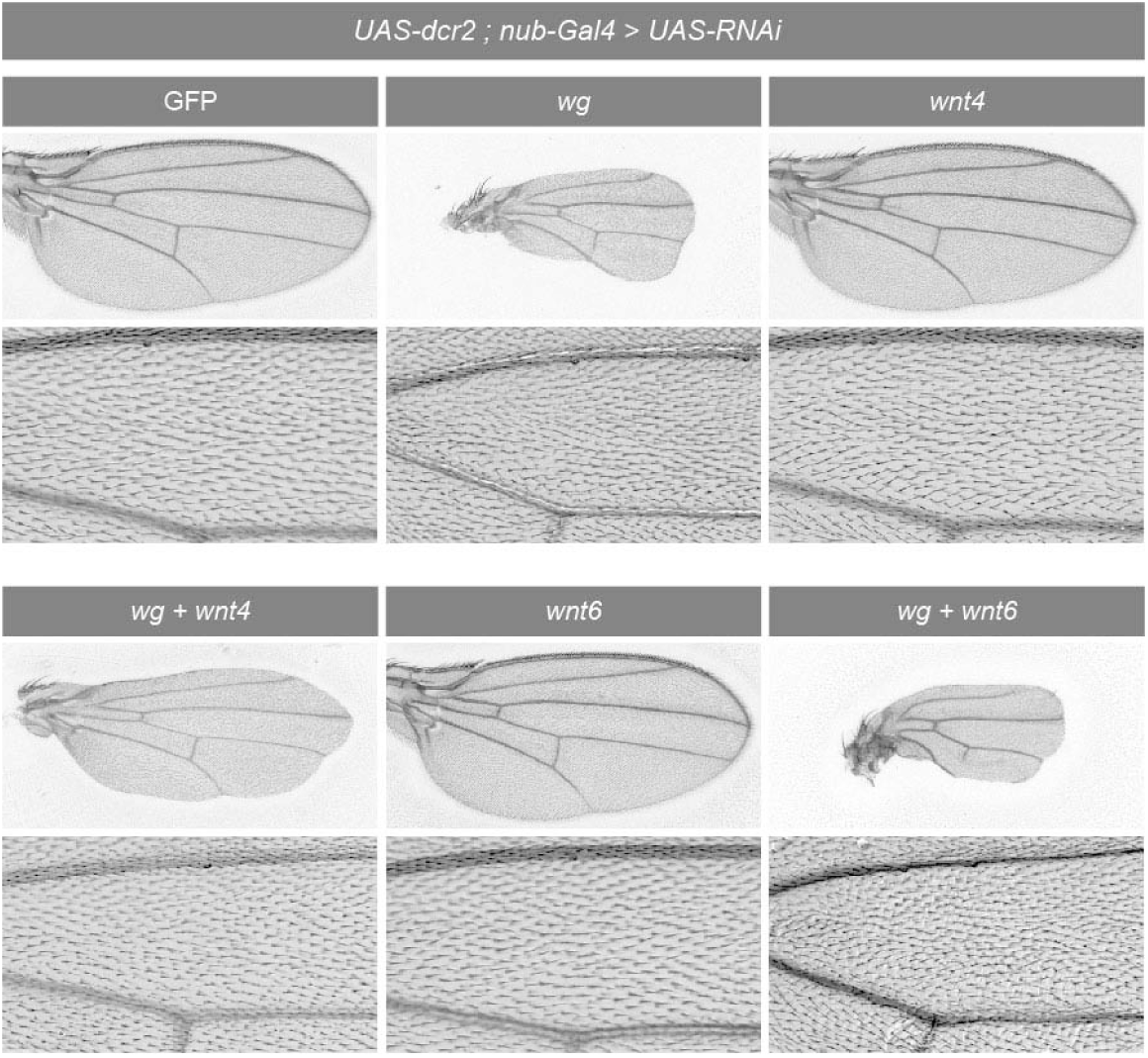
Pairwise double RNAi of *wg, wnt4*, and *wnt6* does not disrupt PCP patterning in the wing. *UAS:RNAi* constructs targeting the indicated genes were co-expressed with *UAS:dcr2* in the developing wing using *nub-Gal4*. Overall wing morphology is shown in top rows, wing hair orientation in the L3-L4 intervein region is shown in magnification in bottom rows. Images show representative images approximately 20 wings analyzed per genotype.

Frizzled-dependent PCP phenotypes can also be studied in the adult notum, by the orientation of bristles (**Figure S7A**). We thus used pairwise RNAi to knock-down *wg* and *wnt4* or *wg* and *wnt6* in the adult notum using *pnr-*Gal4. Consistent with our observations in the adult wing, we observed no discernable PCP phenotypes in the adult notum in any of these conditions (**Figure S7**).

## Discussion

In this study, we performed loss-of-function studies in the wing and notum using multiplexed somatic CRISPR and double RNAi to systematically target Wnt ligands, and in no combination did we observe PCP phenotypes. These results suggest that Wnt ligands are not, whether singly or redundantly with one another, necessary to provide an instructive global cue to the core PCP pathway. If there is a secreted “Factor X” molecule that provides tissue-scale information, our results suggest that it is not a Wnt ligand or combination of multiple Wnt ligands. Previous experiments have suggested that this hypothetical factor is not a member of the Notch, EGF, FGF, or Dpp signaling pathways (Lawrence et al., 2002).

Our results do not rule out the possibility that Wnt ligands could play a redundant role with an additional global cue or cues. For example, a number of studies have suggested that signaling via Fat, Daschous, and Four-jointed may provide large-scale positional information that is interpreted and refined by the activity of core PCP components within and between neighboring cells (reviewed in Butler and Wallingford, 2017; Goodrich and Strutt, 2011). The mechanisms that couple local and global PCP thus remain incompletely understood.

Our results imply that the ability of ectopically expressed Wg and Wnt4 to reorient the PCP of surrounding cells is not reflective of such a role in wildtype flies. One possible explanation for these results is that feedback mechanisms between Wnt ligands and Fz proteins may account for the ability of ectopic Wnt ligands to disrupt wildtype Fz function (Cadigan et al., 1998; Chaudhary et al., 2019), irrespective of the physiological role for Wnts under normal conditions.

During the preparation of this manuscript, a preprint from Yu *et al*. (2020) reported related data and similar conclusions using several independent approaches. Specifically, Yu *et al*. (2020) found that normal PCP patterning occurs in the wings of double mutant flies expressing only membrane-tethered Wg and homozygous for mutations for *wnt4*, as well as in quintuple mutant lacking *wnt2, wnt4, wnt6, wnt10*, and expressing solely membrane-tethered Wg. They also report that *evi/wntless* loss-of-function does not interfere with PCP, nor does driving apoptosis in all *wg*-expressing cells in the developing disc by over-expressing *hid* and *rpr*.

One experimental difference between our studies and those reported by Yu *et al*. (2020) is that the latter employ a membrane-tethered Wg mutant, rather than a knockout or knock-down approach, when addressing Wg function. These mutants lack a gradient of secreted Wg, yet unlike *wg* loss-of-function mutants, display nearly normal patterning and development (Alexandre et al., 2014). Subsequent studies have attributed this phenomenon to a combination of *fz2-*dependent feedback and perdurance of membrane-tethered Wg from earlier stages (Chaudhary et al., 2019). Thus, it is formally possible that membrane-tethered Wg could serve as a global PCP cue in these experiments, even if it not secreted. In the present study, we employ both somatic CRISPR and RNAi to greatly reduce or eliminate *wg* function altogether. We note, for example, that our *wg* RNAi phenotypes resemble those induced by inducing *wg-* expressing cells altogether, and thus likely represent very strong loss-of-function phenotypes, yet we also observed no PCP phenotypes. Taken together, the results presented in the present study are complementary and consistent with those presented by Yu *et al*. (2020), and both studies fail to find evidence that Wnt genes function in PCP in the *Drosophila* wing.

We note that while both Yu *et al*. (2020) and the present study find *wnt4* expression at the wing margin during pupal stages, we did not detect *wnt4* reporter expression in the DV boundary during the L3 larval stage. Although we confirmed our results using three independent knock-in approaches, our results on this point conflict with multiple published studies using *in situ* hybridization (Chen et al., 2008; Gieseler et al., 2001; Harris et al., 2016) as well as the GFP knock-in described by (Yu et al., 2020). It is possible that *wnt4* expression levels are very low, or *wnt4* expression is partially regulated by DNA sequences disrupted by the three knock-in constructs utilized in this study.

### A note on somatic CRISPR versus RNAi

A number of studies have demonstrated that *in vivo* somatic CRISPR is an effective and scalable approach for loss-of-function studies (Poe et al., 2018; Port and Bullock, 2016; Port et al., 2020), and the results in the present study further demonstrate this approach can also be applied to efficiently perform multiplexed somatic knock-outs in a tissue-specific manner. Our results also illustrate a known caveat of the approach which may be especially relevant for secreted factors: somatic CRISPR leads to mosaic tissues that contain a mixture of wildtype and mutant cells. While the proportion of mutant cells can be increased by using multiple sgRNAs (Port and Bullock, 2016), it is unlikely to ever reach 100%, and variability between individuals is likely to remain an issue. For secreted proteins such as Wg, even small numbers of remaining cells that secrete wildtype protein may affect the interpretation of results. *In vivo* RNAi, on the other hand, tends to be far more uniform within and between individuals, and in those cases where particularly effective reagents are available like those used in the present study (compare *wg* loss-of-function phenotypes in **Figure 2** and **Figure 3**), this can result in stronger tissue-level phenotypes than somatic CRISPR. Thus, the advent of somatic CRISPR does not obviate the power of *in vivo* RNAi for functional studies.

## Methods

*Fly work. Drosophila melanogaster* stocks were maintained and crossed on standard laboratory cornmeal food, and all experiments were conducted at 25ºC. The following previously described stocks were used:

*nub-Gal4*; *UAS:Cas9*.*P2* (BL67086)

*UAS:dcr2*; *nub-Gal4* (BL25754)

*pnr-Gal4 / TM3* (Perrimon lab stock)

*elav-Gal4*; *UAS:Cas9*.*P2* (BL67073)

split-Gal4 “p65 tester line”: *tub::Gal4DBD, UAS:EGFP* (BL60298)

split-gal4 “Gal4DBD tester line*”: tub::VP16[AD], UAS:EYFP* (BL60294)

*wnt4* CRIMIC “trojan Gal4” (BL67449)

GTRACE: *UAS-RedStinger, UAS-Flp1*.*D, Ubi (FRT*.*Stop)Stinger/CyO* (BL28280)

GFP RNAi: HMS00314

*pCFD6-wntless*^*2X*^ – gift of F. Port (Port and Bullock, 2016)

*wg* RNAi – dsRNA, VDRC (KK104579)

*wnt4* RNAi – dsRNA, VALIUM10 vector (BL29442)

*wnt6* RNAi – dsRNA, VDRC (GD26669)

*sgRNAs construction*. For each Wnt ligand, five candidate sgRNAs were designed using the *Drosophila Resource Screening Center Find CRISPR v2* online tool (http://www.flyrnai.org/crispr2/). These five candidate sgRNAs were cloned into U6:3-expression vector pCFD3 (Port et al., 2014), and tested for cutting efficiency in S2R+ cells engineered to express Cas9 (Viswanatha et al., 2018), followed by T7 endonuclease activity three days after transfection. Cutting efficiency was estimated by quantifying band intensity via ImageJ, and the top two scoring sgRNAs were selected for usage in this study. *In vivo* guides were cloned into pCFD6, which expresses multiple sgRNAs under UASt control, each flanked by tRNA sequences (Port and Bullock, 2016). The sgRNA sequences used in this study are shown in **Table S1**. For single gene knock-out experiments, two sgRNAs were cloned into the pCFD6 backbone and inserted into the attP2 landing site on the third chromosome, and for double gene knock-out experiments, four sgRNAs were cloned into pCFD6, two sgRNAs per target gene, and inserted into the attP40 site on the second chromosome, using standard PhiC31 recombination.

*Construction of pCFD4*^*FLPOUT*^ (used in **Figure S6**). For pairwise *frizzled* knock-out experiments, we created a modified pCFD4 variant (Port et al., 2014) based on the CoinFLP approach described in (Bosch et al., 2015). In this two-guide sgRNA vector, each sgRNA is flanked by a distinct fluorescent reporter, and interlocking FRT/FRT3 sites, for a final orientation of *FRT-3xP3-GFP-U6:1-sgRNA1-FRT3-FRT-U6:3-sgRNA2-3xP3-mCherry-FRT3*. This fragment was synthesized by GENEWIZ, Inc. (South Plainfield, NJ) and subcloned into the pCFD4 backbone using Gibson cloning.

We reasoned that this plasmid would allow us to create a single transgenic fly line expressing two guides, and to subsequently generate each of the two single sgRNA components by crossing this “parent” line to a source of *hsFlp* and screening the offspring for those that underwent each of the two possible recombination events (FRT vs. FRT3), which would express either GFP and mCherry. We validated that this construct successfully expressed two sgRNAs targeting *ebony* and *forked*, which both produce visible adult phenotypes (not shown). We also validated that the FLP-FRT procedure could be used to successfully generate two stable single-sgRNA lines from one “parent” double-sgRNA line. This construct may be of interest for those wishing to screen large numbers of pairs of genes, and subsequently perform single gene studies on a smaller subset of those genes.

*T7 endonuclease assay*. To test for *in vivo* target cleavage of Wnt genes, pCFD6 males were crossed to unmated *elav-Gal4; UAS:Cas9*.*P2* females, and the adult heads of their F1 offspring were collected for genomic DNA. ∼5-10 heads were collected on dry ice and genomic DNA was extracted by physically homogenization in DNA extraction buffer (10mM Tris-Cl pH 8.2, 1mM EDTA, 25mM NaCL, 200 µg/mL Proteinase K), followed by incubation at 37°C for 20-30 minutes, then boiling at 98°C for ∼90 seconds. 1 µL of genomic DNA was used as template in a 20 µL PCR reaction using primers described in **Table S2** to amplify regions surrounding the sgRNA target site. PCP products were used as template for T7 endonuclease assay (New England BioLabs), following manufacturer instructions, and digested products were visualized on a 2% agarose gel.

*Generation of split-Gal4 knock-in reporter lines*. To create split-Gal4 donor vectors (both p65 and Gal4DBD), p65 and Gal4DBD amplified from pBPp65ADZpUw (Addgene 26234) and pBPp65ADZpUw (Addgene 26233), respectively, and subcloned into pHD-dsRed-attP (Addgene 51019)(Gratz et al., 2014; Pfeiffer et al., 2010). For each Wnt gene, homology arms of approximately 1 kb flanking the sgRNA were designed to create an in-frame fusion between the first exon of the target gene and the T2A sequence of the donor construct, and were cloned into donor vector using Gibson cloning.

Donor constructs and sgRNAs (in pCFD3) were co-injected into *yv;; nanos:Cas9* flies. Injected offspring were backcrossed to balancer stocks on the X, II, or III chromosome depending on the target gene, and screened for RFP+ eyes. Multiple transformants were recovered for both p65 and Gal4DBD knock-ins of each Wnt gene, with the exception of *wnt4* and *wnt10*, for which only p65 and Gal4DBD lines were recovered, respectively. Expression patterns were ascertained by crossing to ubiquitous split-Gal4 “tester lines” (BL60298 and BL60294), which ubiquitously express either VP16 or Gal4DBD together with a UAS-driven reporter. For every Wnt-family knock-in created, multiple transformants gave indistinguishable expression pattern, regardless of whether p65 or Gal4DBD was knocked-in. PCR analyses confirmed that each line was inserted in the predicted location, with the exception of *wntD*, which we were unable to conclusively validate via PCR. We note that all *wntD* knock-in lines, independently generated, and both with p65 and Gal4DBD, recapitulate indistinguishable expression patterns from one another.

*Generation of additional Wnt4 reporter lines*. An additional *wnt4* reporter line was generated using a “universal donor” homology-independent knock-in strategy (Bosch et al., 2019). Briefly, a universal T2A-Gal4, 3xP3-dsRed donor was injected into *yv*;; *nos::Cas9* embryos along with two sgRNAs: on to linearize the donor construct and one to target in the last exon of *wnt4*. These injected G0 flies were crossed to a *w*; *Sp/CyO*; *UAS-GFP* balancer line and screened for RFP and GFP expression. Of 28 fertile G0s six were RFP+, five were GFP-negative, indicating they were likely out-of-frame and/or inserted at the wrong location. One of the six RFP+ lines was GFP+, and had an expression pattern indistinguishable from the two other *wnt4* lines tested in this study.

*Antibody staining and imaging*. Tissues were dissected in cold PBS, fixed for 20-30 minutes in 4% paraformaldehyde in PBS, and then stained following standard immunhistochemical protocols. Tissues were stained with rabbit anti-GFP conjugated to Alexa 488 (Molecular Probes, 1:400) and DAPI (1:1000), and imaged using either a Zeiss LSM 780 or an Olympus IX83 confocal microscope, both part of the Microscopy Resources of the North Quad (MicRoN) facility at Harvard Medical School. Maximum intensity projections are shown for all tissues except the proventriculus, for which single z-slices are also shown. Adult wings were mounted in a 1:1 mixture of Permount (Fisher) and xylenes, and imaged using brightfield optics on a Zeiss Axioskop 2. Nota were imaged using a Zeiss AxioZoom, and z-stacks were compiled into a single image using Helicon Focus software.

## Acknowledgements

We thank Fillip Port for providing pCFD6-*evi* flies, Marco Milan and Lara Barrios for providing combined RNAi lines targeting *wg, wnt4*, and *wnt6*, Rich Binari for assistance with fly work, Justin Bosch for assistance designing pCFD4^FLPOUT^, and Pedro Saavedra, Stephanie Mohr, Sudhir Tattikota, and members of the Perrimon Lab for critical feedback. BEC received funding from the Charles A. King Postdoctoral Research Fellowship program, and the Perrimon Lab received funding from NIH ORIP R24 OD26435 and HHMI.

## Author contributions

B.E.C. designed and performed experiments, generated reagents, and wrote the manuscript. T.C. performed gut imaging experiments. E.V. generated reagents. N.P. designed experiments and edited the manuscript.

**Supplementary Note 1**. Using the split-Gal4 lines described in this study, we examined Wnt ligand expression in the adult ovary, larval CNS, and two stem cell populations in the adult gut **(Figure S3, S4**). In the adult ovary, our split-Gal4 reporters demonstrated that *wg, wnt2, wnt4*, and *wnt6* are co-expressed in a partially overlapping pattern in the female germarium, matching previous descriptions (Waghmare and Page-McCaw, 2018). In addition, we found that *wnt5* is also expressed in this tissue, in a pattern similar to *wnt6* (**Figure S3A**), which to our knowledge has not been previously reported.

We find that a similar subset of Wnt ligands are co-expressed in and surrounding adult stem cells located in the adult gut: those in the cardia (located at the foregut-midgut junction), and those at the midgut-hindgut juncture (**Figure S4**) (Singh et al., 2011; Tian et al., 2016). We observed that four of these same Wnt genes expressed in the germarium were in the cardia (*wg, wnt2, wnt4*, and *wnt6*), and three (*wg, wnt4*, and *wnt6*) are co-expressed in the stem cells at the midgut-hindgut juncture. These observations add to our understanding of which ligands are responsible for Wnt signaling observed in these regions of the gut (Tian et al., 2016, 2018).

We also examined Wnt gene expression in the larval nervous system. *wg, wnt4*, and *wnt5* are known to have roles in the developing nervous system (Inaki et al., 2007; Packard et al., 2002; Yoshikawa et al., 2003), and we find that in fact all Wnts are expressed in the larval nervous system. This includes *wnt10*, whose expression pattern, to our knowledge, has not been previously described (**Figure S3**).

---

**Fig S1.**
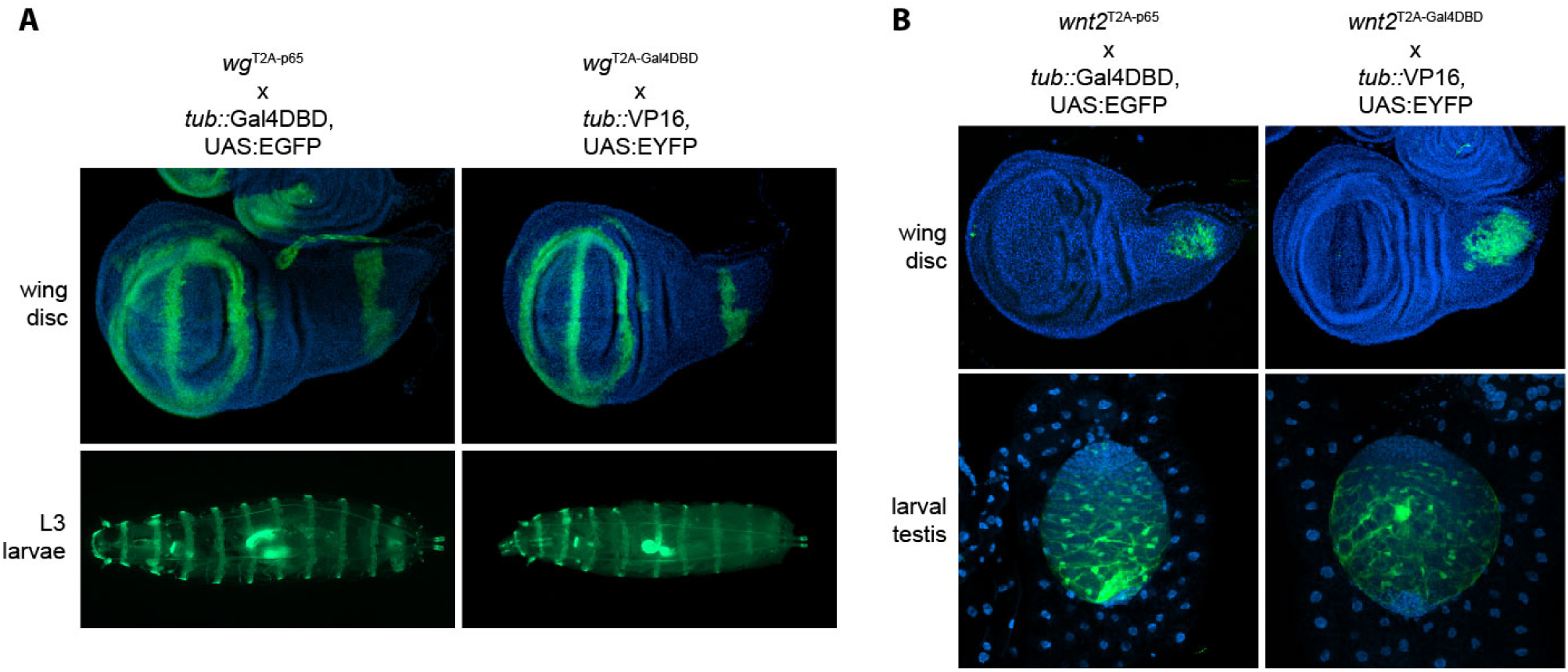
Pilot characteriziation of split-gal4 knock-in reporters for *wg* and *wnt2*. Split-Gal4 expression patterns revealed by crossing to a ubiquitously-expressed split-Gal4 “tester” line. **(A)** split-Gal4 reporter expression in the canonical Wg expression pattern in L3 larval wing disc (anterior is down), and in L3 larvae (anterior is left), revealed for both p65 and Gal4DBD knock-ins. (A) *wnt2* is expressed in the presumptive notum at the base of the wing disc, and in the larval testis, consistent with its described roles in thoracic muscle and testis development. UAS:EGFP/EYFP expression is shown in green, and DAPI is shown in blue.

**Fig S2.**
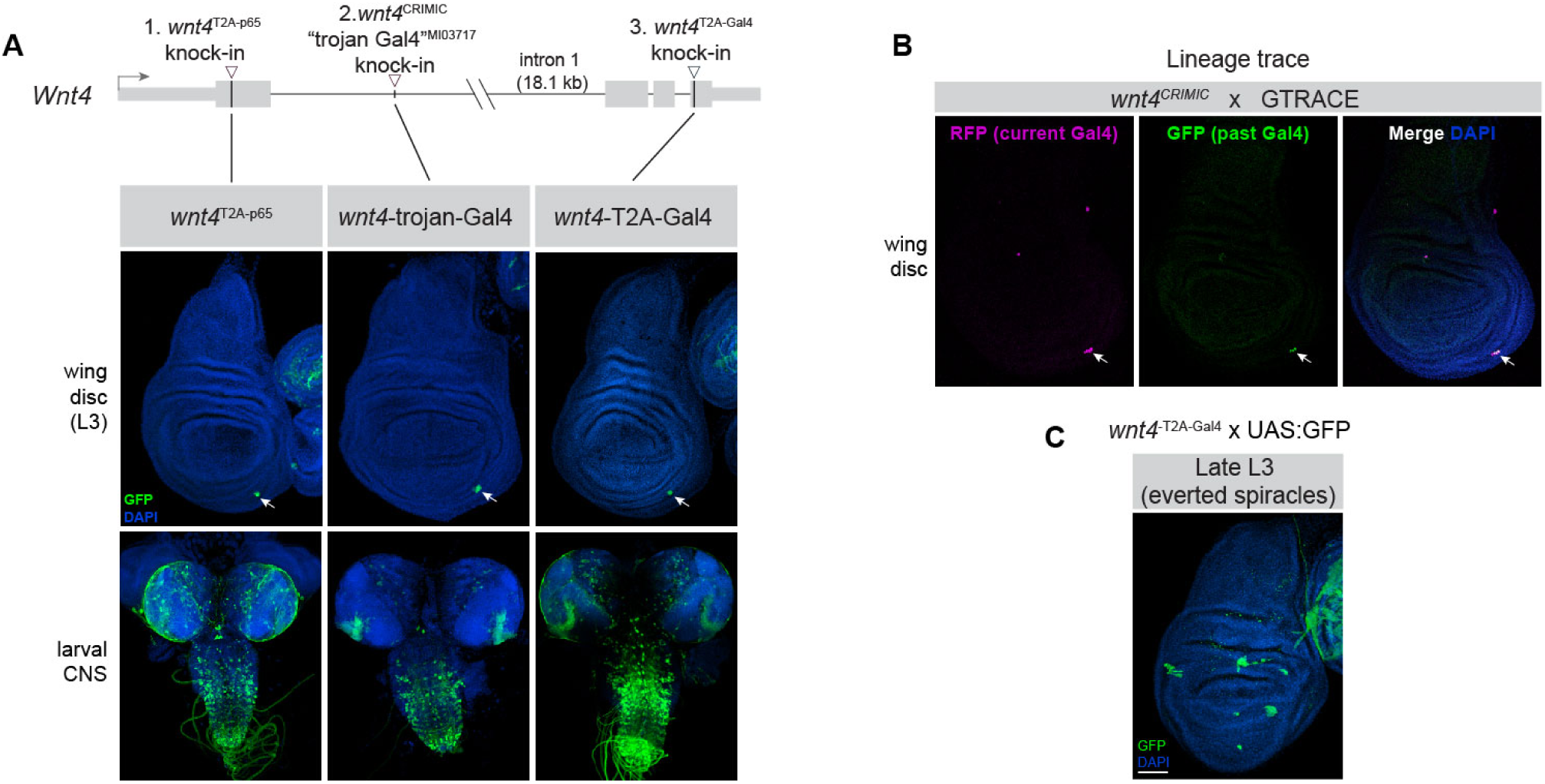
Three independent *wnt4* knock-in reporters are largely consistent with one another and suggest low or absent expression in the L3 wing margin. (A) Schematic of three independent knock-in reporters in the *wnt4* locus, in the first exon, first intron, and last exon, and their expression pattern in the L3 wing disc and in the larval CNS. (B) Lineage tracing of a “trojan Gal4” Wnt4 insertion using GTRACE indicates that *wnt4* is not expressed during wing development prior to L3 (C) *wnt4* expression expands in late larval stage (everted spiracles stage). White arrows in (A) and (B) indicate a consistently labeled population of cells on the antero-ventral margin of the wing pouch.

**Fig S3.**
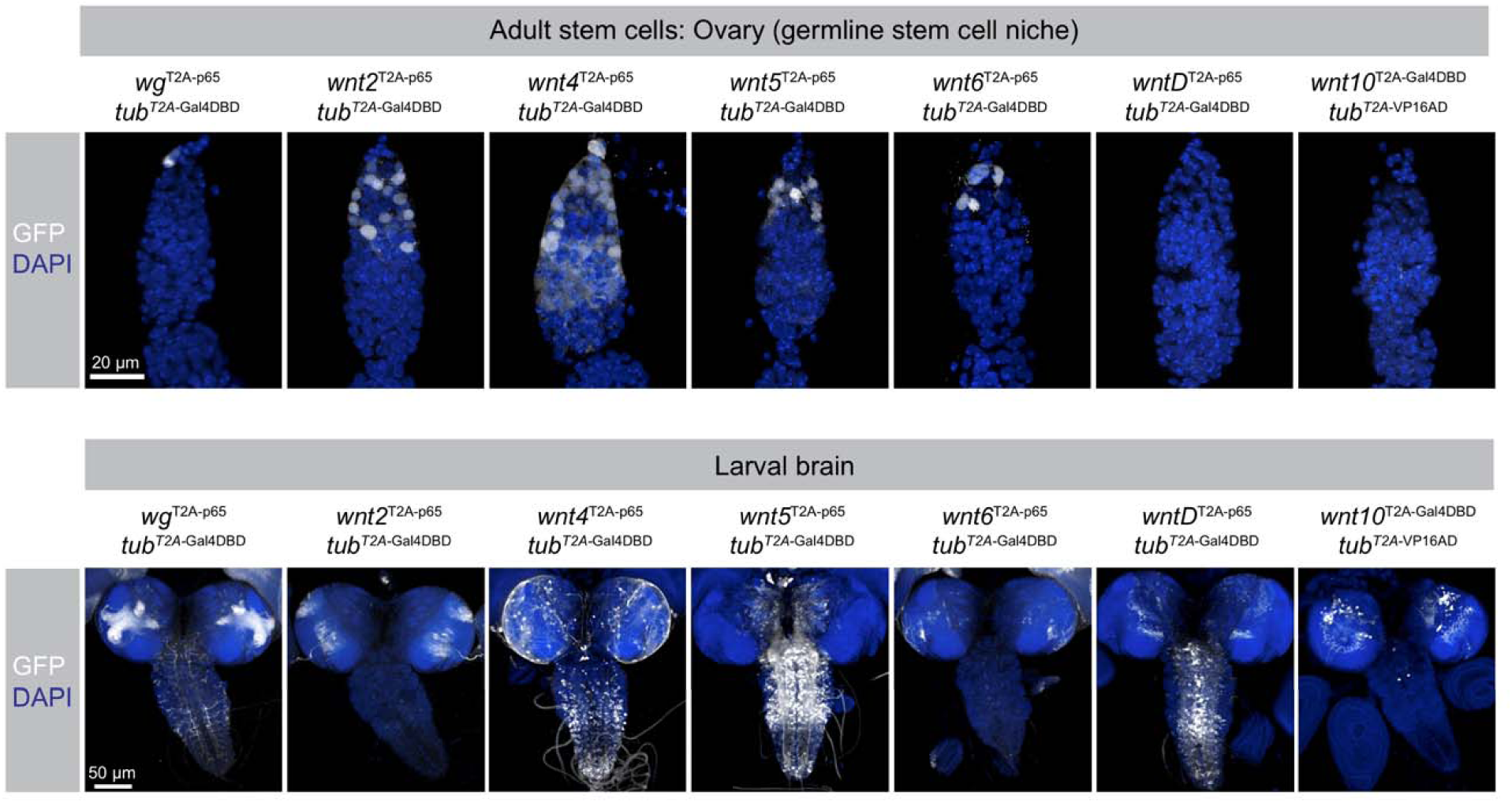
Additional characterization of split-Gal4 Wnt reporters in larval and adult tissues. Wnt split-gal4 reporter expression in (A) the ovarian germline stem cells and surrounding tissues of the adult germarium, and (B) the larval CNS.

**Fig S4.**
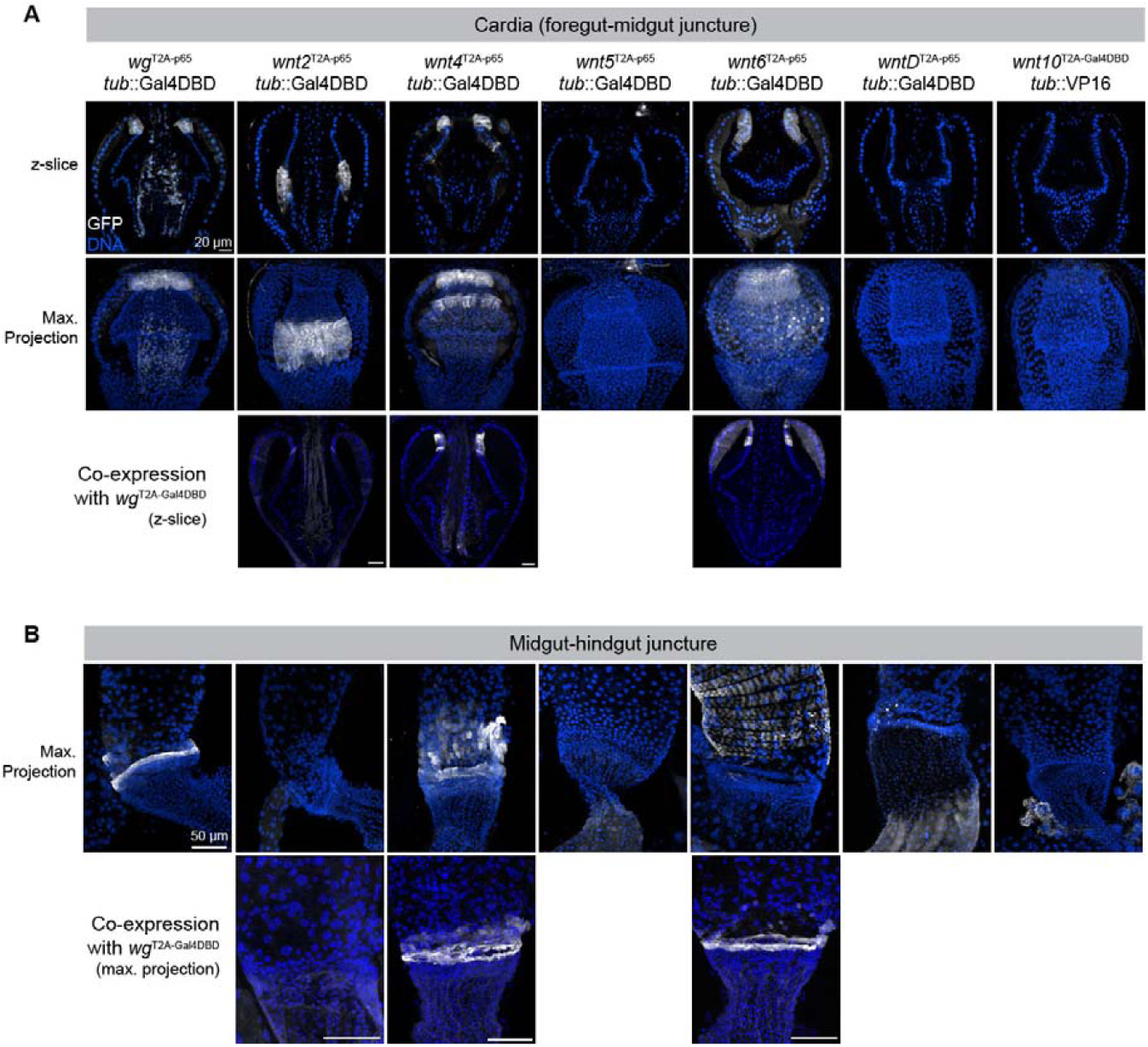
Multiple Wnt genes are expressed in populations of adult stem cells located at regional junctions of the gut. (A) Wnt split-gal4 reporter expression in stem cells and surrounding tissue of the adult cardia (foregut-midgut juncture) and (B) in a band of cells at the midgut-hindgut junction.

**Fig S5.**
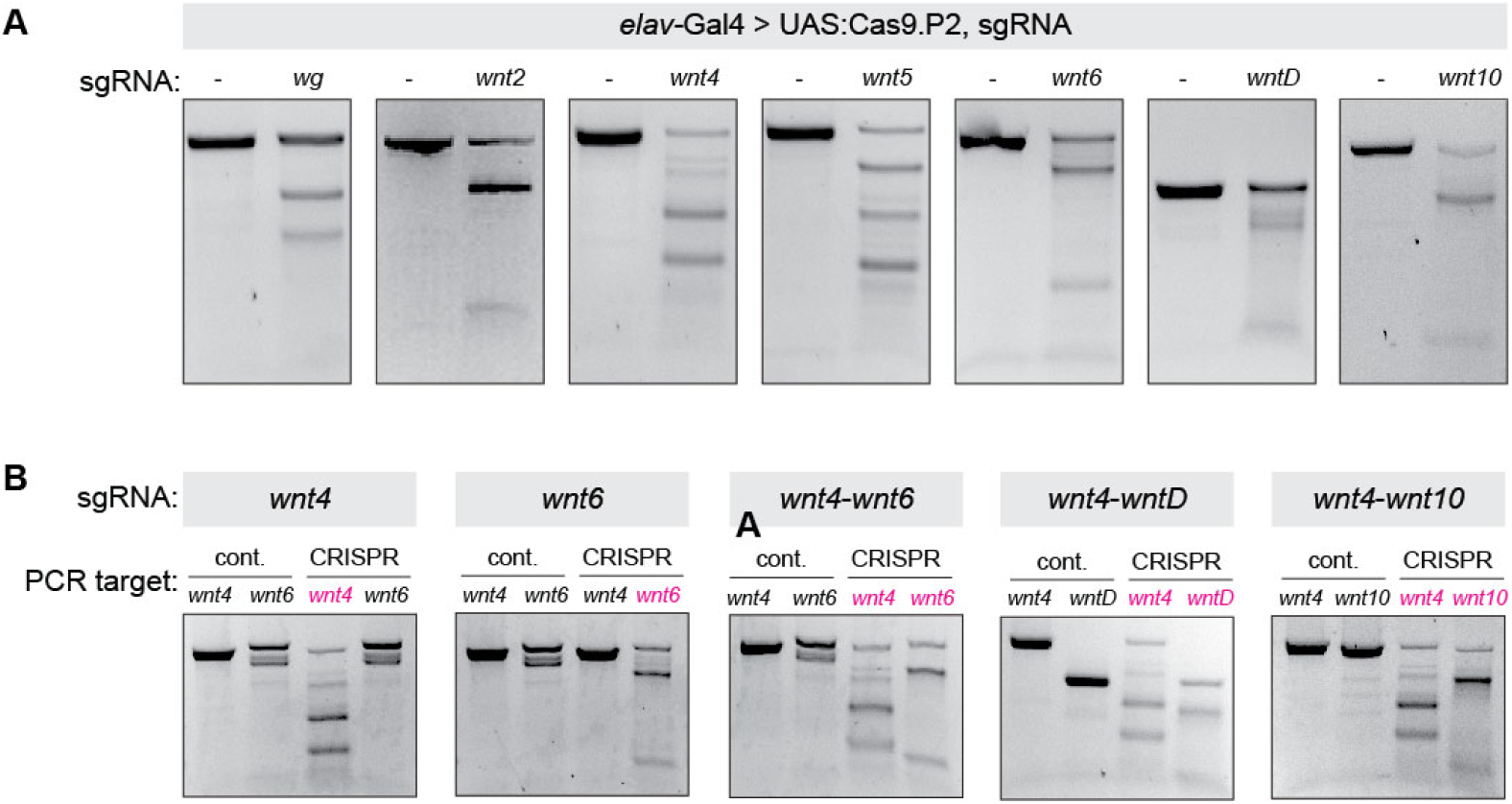
Effective cleavage of Wnt genes, both singly and in pairs, using multiplexed *in vivo* somatic CRISPR. Two sgRNAS per target Wnt gene were cloned into the pCFD6 backbone, which expresses multiple sgRNAs under UAS^t^ control, each separated by tRNAs. *UAS:Cas9*.*P2* and *UAS:sgRNAs* were expressed in the adult nervous system using *elav-Gal4*, and T7 assays were conducted on adult heads. **(A)** *In vivo* CRISPR-mediated cleavage of each Wnt gene visualized by T7 endonuclease activity. **(B)** Simultaneous *in vivo* CRISPR cleavage of pairs of Wnt genes. Target gene is indicated in magenta. Note that each gene is targeted by two sgRNAs, and that in some cases the PCR-amplified fragment used for the T7 endonuclease assay only includes one such target site.

**Fig S6.**
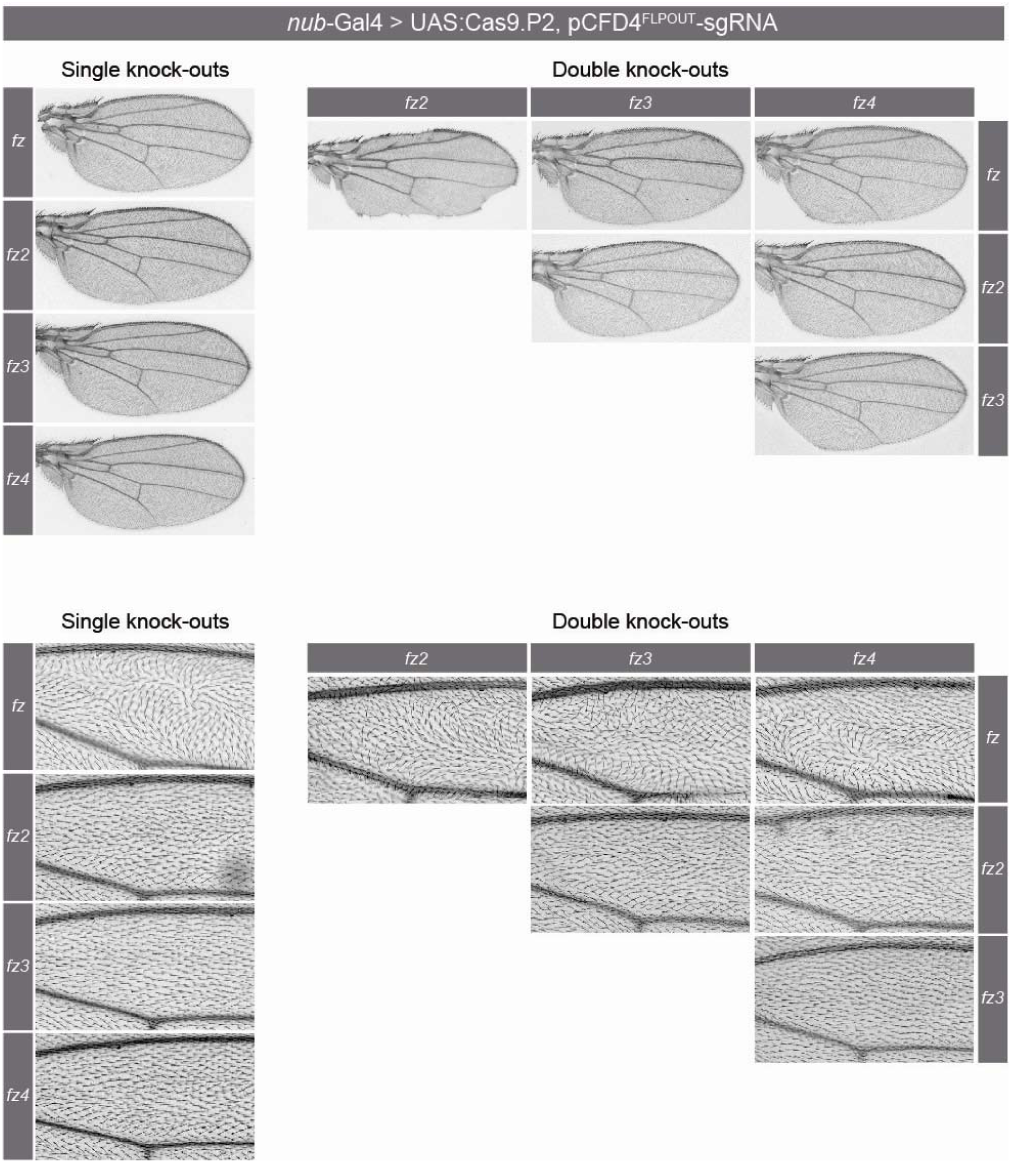
Somatic CRISPR of *fz*, but not *fz2, fz3*, or *fz4* nor any combination thereof, produces planar cell polarity defects in the adult wing. Each of the four *fz* paralogs was targeted (one sgRNA per target gene, in a modified pCFD4 backbone [see Methods]) in the wing using *nub-Gal4 > UAS:Cas9*.*P2*, either singly or in each pairwise combination, and morphology and PCP was assayed in the adult wing. Top panels show whole phenotype (note that wing margin defects are specific to double knock-out of *fz* + *fz2*), and bottom panels show higher magnification of wing hair orientation in the of the L2-L3 intervein region. PCP phenotypes were exclusively observed when *fz* was targeted.

**Fig S7.**
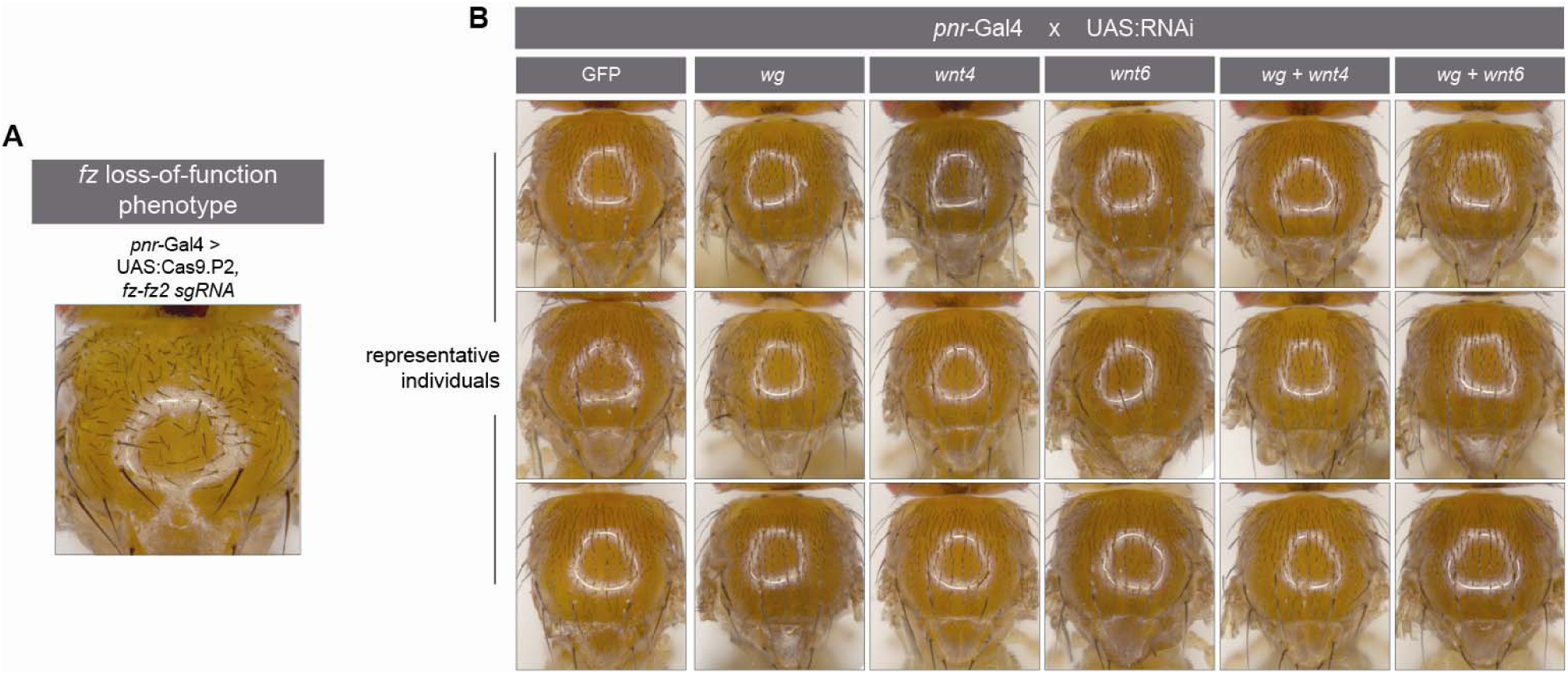
Pairwise double RNAi against *wg* in combination with *wnt4* or *wnt6* in the notum does not disrupt PCP patterning. **(A)** Example of a characteristic PCP phenotype (misoriented bristles) in the adult notum caused *fz* loss-of-function (*pnr-Gal4 > UAS:Cas9*.*P2, sgRNA-fz-fz2*). **(B)** *UAS:RNAi* constructs targeting the indicated genes were expressed in the developing notum using *pnr-Gal4*, and PCP was analyzed by observing bristle polarity.

**Table S1.**
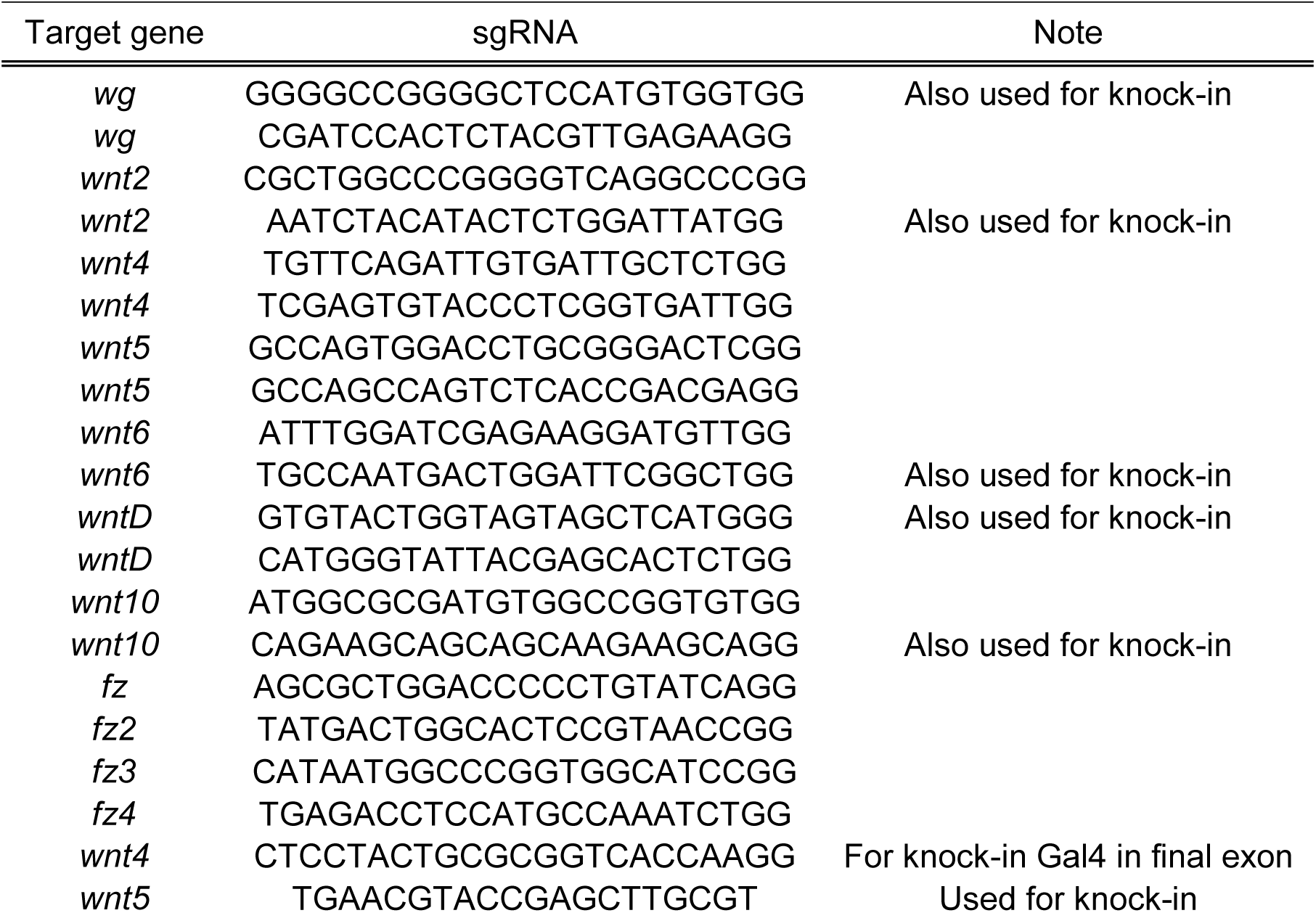
sgRNA sequences used in this study.

**Table S2.**
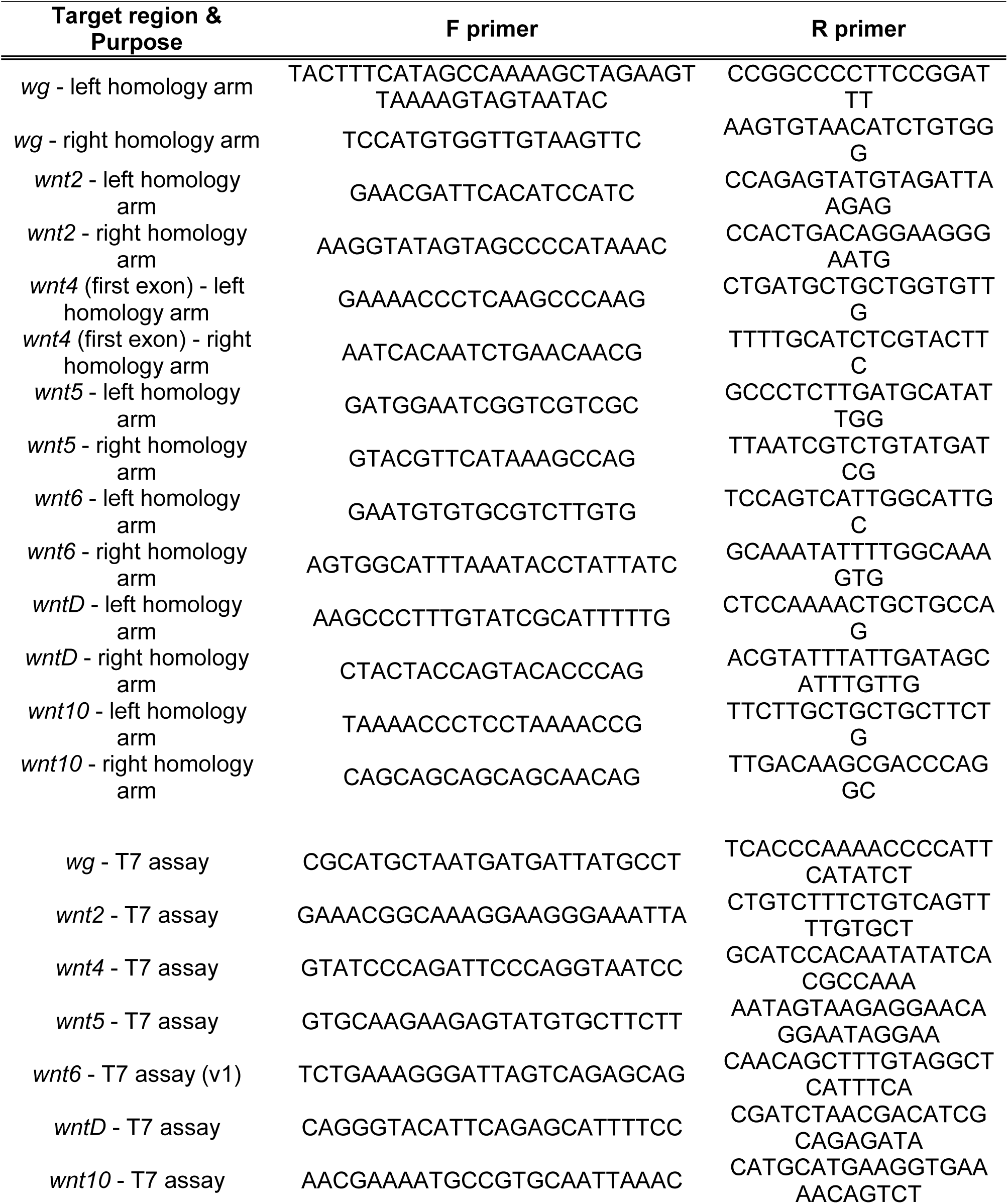
PCR primers used in this study.

